# Corrigendum and follow-up: Whole genome sequencing of multiple CRISPR-edited mouse lines suggests no excess mutations

**DOI:** 10.1101/154450

**Authors:** Kellie A. Schaefer, Benjamin W Darbro, Diana F. Colgan, Stephen H. Tsang, Alexander G. Bassuk, Vinit B. Mahajan

**Author notes:** Corresponding Author: Vinit B. Mahajan M.D., Ph.D., Byers Eye Institute, Department of Ophthalmology, Stanford University, Palo Alto, CA, 94304 USA. Alexander G. Bassuk, M.D., Ph.D., Department of Pediatrics, The University of Iowa, Iowa City, IA, 52242.

## Abstract

Our previous publication suggested CRISPR-Cas9 editing at the zygotic stage might unexpectedly introduce a multitude of subtle but unintended mutations, an interpretation that not surprisingly raised numerous questions. The key issue is that since parental lines were not available, might the reported variants have been inherited? To expand upon the limited available whole genome data on whether CRISPR-edited mice show more genetic variation, whole-genome sequencing was performed on two other mouse lines that had undergone a CRISPR-editing procedure. Again, parents were not available for either the Capn5 nor Fblim1 CRISPR-edited mouse lines, so strain controls were examined. Additionally, we also include verification of variants detected in the initial mouse line. Taken together, these whole-genome-sequencing-level results support the idea that in specific cases, CRISPR-Cas9 editing can precisely edit the genome at the organismal level and may not introduce numerous, unintended, off-target mutations.

## Introduction

Our previous publication suggested CRISPR-Cas9 editing at the zygotic stage might unexpectedly introduce a multitude of subtle but unintended mutations, an interpretation that not surprisingly raised numerous questions. The key issue is that since parental lines were not available, might the reported variants have been inherited?^1^ To expand upon the limited available whole genome data on whether CRISPR-edited mice show more genetic variation, whole-genome sequencing was performed on two other mouse lines that had undergone a CRISPR-editing procedure. The first line was created using the C57Bl6/J background and introduced a non-synonymous point mutation in the *Capn5* gene. The second mouse line was created using the C57Bl/6NCrl background and introduced a frameshift mutation in the *Fblim1* gene (see Methods for details). Again, parents were not available for either the *Capn5* nor *Fblim1* CRISPR-edited mouse lines, so strain controls were examined.

## Methods

### Mice

The Institutional Animal Care and Use Committee (IACUC) approved all experiments. Rodents were used in accordance with the Statement for the Use of Animals in Ophthalmic and Vision Research of the Association for Research in Vision and Ophthalmology, as well as the Policy for the Use of Animals in Neuroscience Research of the Society for Neuroscience.

C57BL/6J CAPN5-R243L mice were designed and produced by Applied StemCell (Milpitas, CA). Embryos were injected through a cytoplasmic route with an ssODN donor, gRNA and spCas9. Two surrogate CD1 mice were implanted with embryos that passed quality screening. Pups were screened for the desired point mutation by PCR and sequencing before being set up as breeders. Tail tissues of F0 and F1 mice were also screened by PCR and sequencing. gRNA and spCas9 were cloned into the same pBT-U6-Cas9-2A-GFP plasmid.

gRNA sequence: 5’-AGCTGACATGGAGGCCCGCCTGG-3’

ssODN sequence: 5’-

GCTGTGACAGCAGCTGACATGGAGGCCTTGCTGGCATGTGGCCTGGTGAAGGGC CATGCATACGCTGTCACCGATGTGCGCAAGGTGCGCCTGGGCCATGGCCTGCTG-3’

PCR primer Forward: 5’-TGGAGCAGTGAGGATAGACAAGC-3’ PCR primer Reverse: 5’-AAGAATGGGGTGTGTAGATGCCT-3’

The mouse line C57Bl/6N-Fblim1<em5Tcp> was made as part of the NorCOMM2 project at The Centre for Phenogenomics (Toronto, Ontario, Canada), and obtained from the Canadian Mouse Mutant Repository. Zygotes were collected from the oviducts of 0.5 days post coitum (dpc), superovulated and plugged, C57Bl/6NCrl mice. Cumulus masses were treated with hyaluronidase to remove cumulus cells then zygotes were rinsed and maintained in KSOM-AA microdrops under oil at 37°C/6% CO_2_ until injection. A microinjection mix of 20 ng/µL Cas9 mRNA (Life Technologies A25640), 10 ng/µL single-guide RNA and 10 ng/µL single-strand oligo repair template was prepared in loTE microinjection buffer (10 mM Tris, pH 7.6, 0.1 mM EDTA) and injected under continuous flow into the pronuclei of zygotes. Embryos that survived injection were transferred to pseudopregnant ICR recipients for gestation and birth. Born pups were genotyped at 2 weeks of age using DNA isolated from tail biopsies and PCR sequencing (PCR-F: TGGCCTCCTCTGTCTTCATCA, PCR-R: TCCAGAGAGTGAGGCATTGGT). Founders were backcrossed to C57BL/6NCrl mice and progeny were first genotyped by PCR sequencing and allele sequence was then confirmed by subcloning the PCR fragment and sequencing individual clones. Heterozygous sequence verified mice were backcrossed a second time to C57BL/6NCrl prior to intercrossing to produce homozygous mice. Mice were intercrossed three times before being shipped to the University of Iowa. Mice received at the University of Iowa were intercrossed an additional two times. These progeny were sent for whole-genome sequencing.

gRNA sequence: 5’-CTTGTGAAGCCAGACGCGCT-3’

Genotyping at the University of Iowa used additional primers.

PCR primer Forward: 5’-TGTAGCCGTGAGTGAGGAA-3’

PCR primer Reverse: 5’-CCAGGTGTCTTTGTGGGAAG-3’

Mice without CRISPR-mediated correction were used as the functional-deficient control.

### DNA Library Prep and Whole Genome Sequencing

Total DNA was extracted from mouse spleen, as described previously.^2^ DNA quality was determined using Picogreen to determine total mass and Fragment Analyzer to determine DNA integrity. Whole genome sequencing (WGS) libraries were prepared using the Truseq DNA PCR-free Library Preparation Kit in accordance with the manufacturer’s instructions. Briefly, 1ug of DNA was sheared using a Covaris LE220 sonicator (adaptive focused acoustics). DNA fragments underwent bead-based size selection and were subsequently end-repaired, adenylated, and ligated to Illumina sequencing adapters. Final libraries were evaluated using fluorescent-based assays including qPCR with the Universal KAPA Library Quantification Kit and Fragment Analyzer (Advanced Analytics) or BioAnalyzer (Agilent2100). Libraries were sequenced on an Illumina HiSeq 2500 sequencer (v4 chemistry) using 2 × 125bp cycles to a mean coverage of 50x for CRISPR-treated animals and 30x for the control animal at the New York Genome Center.

### Bioinformatic Analysis

Genomic sequencing reads were mapped to the mouse reference genome (GRCm38/mm10) using BWA-mem and subsequent BAM files underwent indel realignment, duplicate marking, and base quality score recalibration as previously described.^1^ Both single nucleotide and indel variants were called simultaneously across all samples using the UnifiedGenotyper and the GRCm38/mm10 reference genome.^3^ Variants were removed from further analysis if the average depth across all samples for the genomic position was less than 30x, the average genotype quality was less than 90, or the variant was present within dbSNP v146 (NCBI. *dbSNP Build 146.* https://www.ncbi.nlm.nih.gov/projects/SNP/snp_summary.cgi?view+summary=view+summary&build_id=146 (2015) and Sherry, S.T. et al. dbSNP: the NCBI database of genetic variation. *Nucleic Acids Res.* 29, 308-311 (2001). Variants were also removed from further analysis if they were part of short tandem repeats or failed any of the GATK’s recommended hard filters (for SNVs, ReadPosRankSum < -8, MQRankSum < - 12.5, Quality by Depth < 2, Strand Bias-Fisher’s > 60, or Mapping Quality < 40, and for indels, ReadPosRankSum < -20, Quality by Depth < 2, Strand Bias-Fisher’s > 200).^3^ Variant filtering was performed using VarSeq (Golden Helix Inc) and variant genotypes were compared using a custom Python script, and the results presented as Venn diagrams created by Venny.^4^

### Topo Cloning and Sanger Sequencing

Mutated regions were amplified using primers (Integrated DNA Technologies), Biolase DNA polymerase (Bioline) and a dNTP mix (New England Biolabs), and subsequently TOPO cloned using the TOPO-TA Cloning Kit (ThermoFisher). Colonies containing the insert were expanded, and the insert was PCR amplified using M13 primers. Crude PCR products were sequenced by the Sanger method (Functional Biosciences).

### High Fidelity Topo Cloning and Sanger Sequencing

DNA fragments were amplified with gene-specific primers, using Phusion High-Fidelity DNA polymerase (M0530, NEB). PCR products were cloned using the Zero Blunt Topo PCR Cloning Kit (K280020, Thermo Fisher Scientific) as per the manufacturer’s instruction. Single colonies were picked the next day and used to inoculate LB media with antibiotics. Bacterial cultures were incubated at 37C for 30 minutes. Bacterial DNA was used as DNA template for PCR amplification with M13 forward and M13 reverse primers using Phusion HF polymerase. Crude PCR products were sequenced using gene-specific primers at Functional Biosciences (Madison, WI).

Primers:

M13 Forward: GTAAAACGACGGCCAGT

M13 Reverse: CAGGAAACAGCTATGAC

Indel chrX:123734765 Forward: CCCTTCACGTTAAACATATTGGA

Indel chrX:123734765 Reverse: TTGACTTACTTTTATATCCAGCCACTT

Indel chr4:66453495 Forward: TTTGGGATGATGGAGGAGAG

Indel chr4:66453495 Reverse: TCATTGTGCCACCAAGAAAC

Indel chr3:88647832 Forward: CAGCCATTTGGAAGAAGCTC

Indel chr3:88647832 Reverse: CCATTCATGCCCTCTTCAGT

Indel chr14:94681371 Forward: ATCCCAGGAAGATTGGCTTT

Indel chr14:94681371 Reverse: GGGAAACGCTTTCAACTATACA

All genome sequences will be uploaded to NCBI upon publication.

## Results

In the previous study, an excess of variants in CRISPR-treated mice when compared to untreated controls was reported. After questions emerged about the interpretation of the data, the variants were verified using TOPO cloning and Sanger sequencing. In addition to detecting the variants reported by WGS, additional variants near Protospacer Adjacent Motif (PAM) sequences in the mutated regions were also detected. Additional analysis suggested inconsistencies with Mendelian inheritance (Supplemental Figures 1-12).

**Figure 1:**
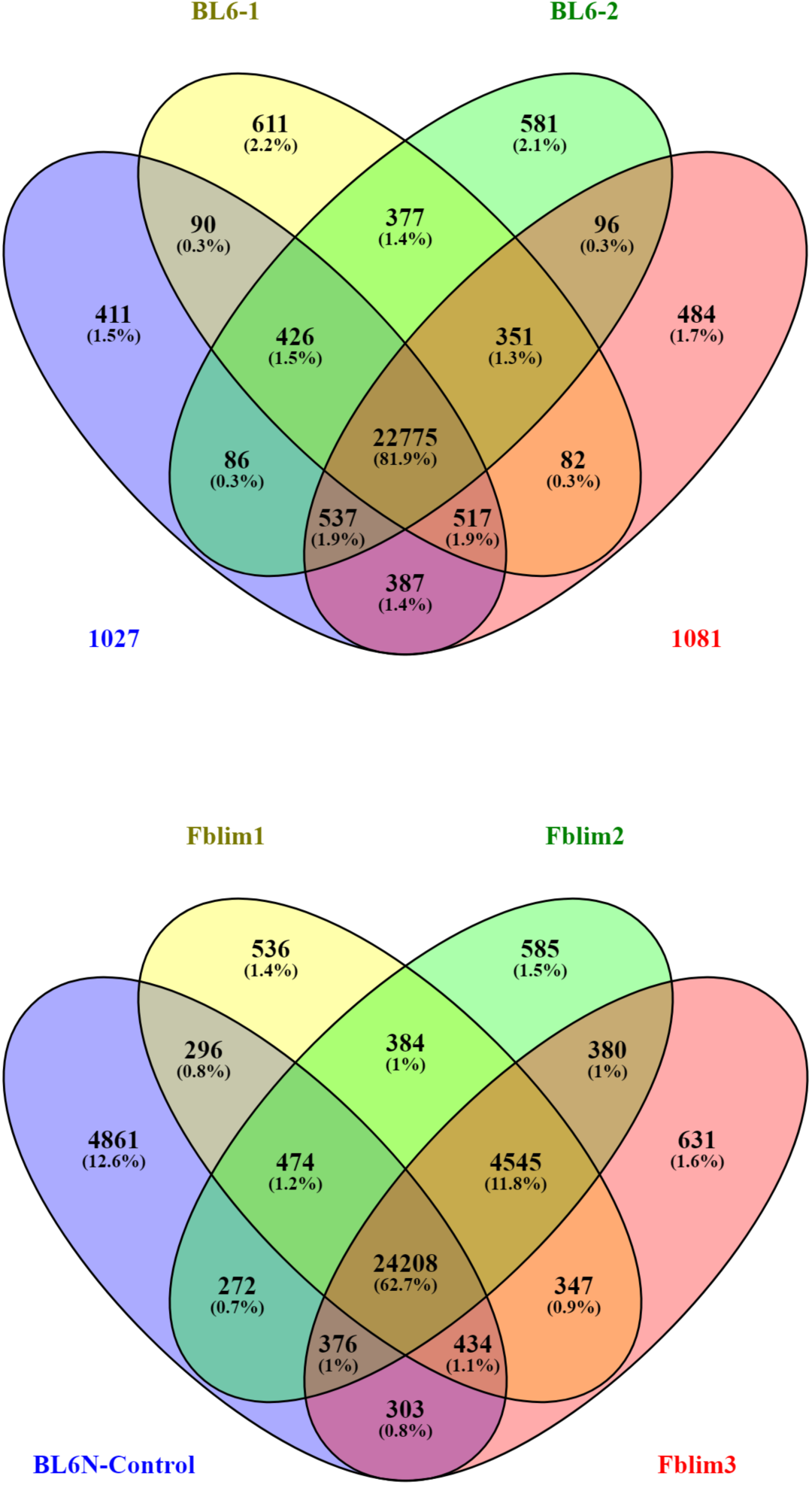
Venn diagrams showing shared and distinct variant genotypes across CRISPR-treated and untreated controls. Samples 1027 and 1081 are Capn5 CRISPR-treated animals and BL6-1 and BL6-2 are the respective strain controls; samples Fblim1, Fblim2, and Fblim3 are the Fblim1 CRISPR-treated animals and BL6N-Control is the respective strain control.

To expand the dataset for analysis, whole-genome sequencing was performed for two groups of four mice. The first group was of the C57Bl6/N background. Three mice had been treated with CRISPR-Cas9 to create a 2bp deletion resulting in a frameshift mutation at A40Sfs*53 in the *Fblim1* gene. The fourth mouse was an untreated control. The other four mice were of the C57Bl6/J background. Two of these mice were treated with CRISPR-Cas9 to create a CGC>TTG mutation in the *CAPN5* gene resulting in the nonsynonymous mutation p.R243L. The other two mice in this group were untreated controls. CRISPR-editing and relatedness of the mice are detailed in Supplemental Table 1. After preliminary bioinformatic analyses similar to those used in the previous study, an excess of unique variants in the CRISPR-edited *Capn5* and *Fblim1* mutant mice were not observed when compared to strain controls. Even after accounting for the 2 generations of backcrossing, this result is different than what was seen in the analysis of the mosaic mice in Schaefer *et al*.

Of note, the previous publication^1^ has a labeling error in Supplemental Figure 3, panel a, mislabeling the location of the top-10, off-target sites predicted by the Benchling software. Initially, when the guide was designed, the mm9 build of the mouse genome was used, but the WGS analysis performed later used the mm10 build, so Figure 3b-d used labels from the mm10 build. Regardless of the build, the sequences in panel 3a are the same, but for consistency, the chromosomal locations and gene names were relabeled using the mm10 build. For clarity, descriptive column titles were added, the first (*Pde6b*) sequence was removed in 3a, and the last five nucleotides of *Pcnt* in panel 3b were corrected (Revised Supplementary Figure 3).

## Discussion

A reasonable concern about unintended effects of CRISPR-Cas9 across the genome and potential associated adverse events if used clinically has been met with limited whole genome sequencing to date. In the current study, dideoxy sequencing was used to verify the variants discussed in our previous letter.^1^ On the other hand, examination of additional CRISPR-treated mice created with a different methodology did not show an excess number of unintended variants potentially introduced by CRISPR-Cas9 gene editing procedure. Each CRISPR-edited mouse was created under different conditions: i.e. pronuclear *vs.* cytoplasmic injection, different gRNAs, different versions of the Cas9 protein expressed from different plasmids, different background strains, and use of backcrossed mice, any or all of which could be associated with different effects of CRISPR-editing in the cell. For example, break-induced replication (BIR) can be triggered by hSpCas9n(D10A) used previously.^5^ Of note, these data suggest that in the absence of parental controls, purchased, inbred mice that are within a few generations of the CRISPR-treated mice may not show significant accumulated genetic drift - their relative value as controls will require evaluation of additional sequence data. Thus the use of same-strain control mice in the original study may not, by itself, explain the excess of variants detected in the previously studied CRISPR-treated mice.^1^

Taken together, these whole-genome-sequencing-level results support the idea that in specific cases, CRISPR-Cas9 editing can precisely edit the genome at the organismal level and may not introduce numerous, unintended, off-target mutations. Although further analysis is needed, along with experiments specifically designed to include parental controls in mice undergoing CRISPR homology directed repair, use of specific gRNAs, updated Cas9 variants, and injection approach may allow for CRISPR-Cas9 gene editing without genome-wide mutations, and suggest a path towards clinically viable CRISPR-based gene editing, including studies to optimize precision and experimentally confirm genome-wide safety.

## Supplemental Figures and Tables

**Supplemental Table 1:** Targeted CRISPR-mutations and relatedness of analyzed mice.

| Mouse ID | Background | Sex | Generation Number | Specific CRISPR-edit |
| --- | --- | --- | --- | --- |
| Fblim1_1 | C57Bl6/N | F | 2 generations backcrossed followed by 5 generations intercrossed | 2bp deletion-p.A40Sfs*53 |
| Fblim1_2 | C57Bl6/N | F | 2 generations backcrossed followed by 5 generations intercrossed | 2bp deletion-p.A40Sfs*53 |
| Fblim1_3 | C57Bl6/N | F | 2 generations backcrossed followed by 5 generations intercrossed | 2bp deletion-p.A40Sfs*53 |
| Control | C57Bl6/N | F | Unknown- ordered from Charles River | Untreated |
| R243L-1027 | C57Bl6/J | M | 2 | R243L CGC>TTG |
| R243L-1081 | C57Bl6/J | M | 2 | R243L CGC>TTG |
| Control 1 | C57Bl6/J | M | Between F226-F236 | Untreated |
| Control 2 | C57Bl6/J | M | Between F226-F236 | Untreated |

**Supplemental Figure 1.**
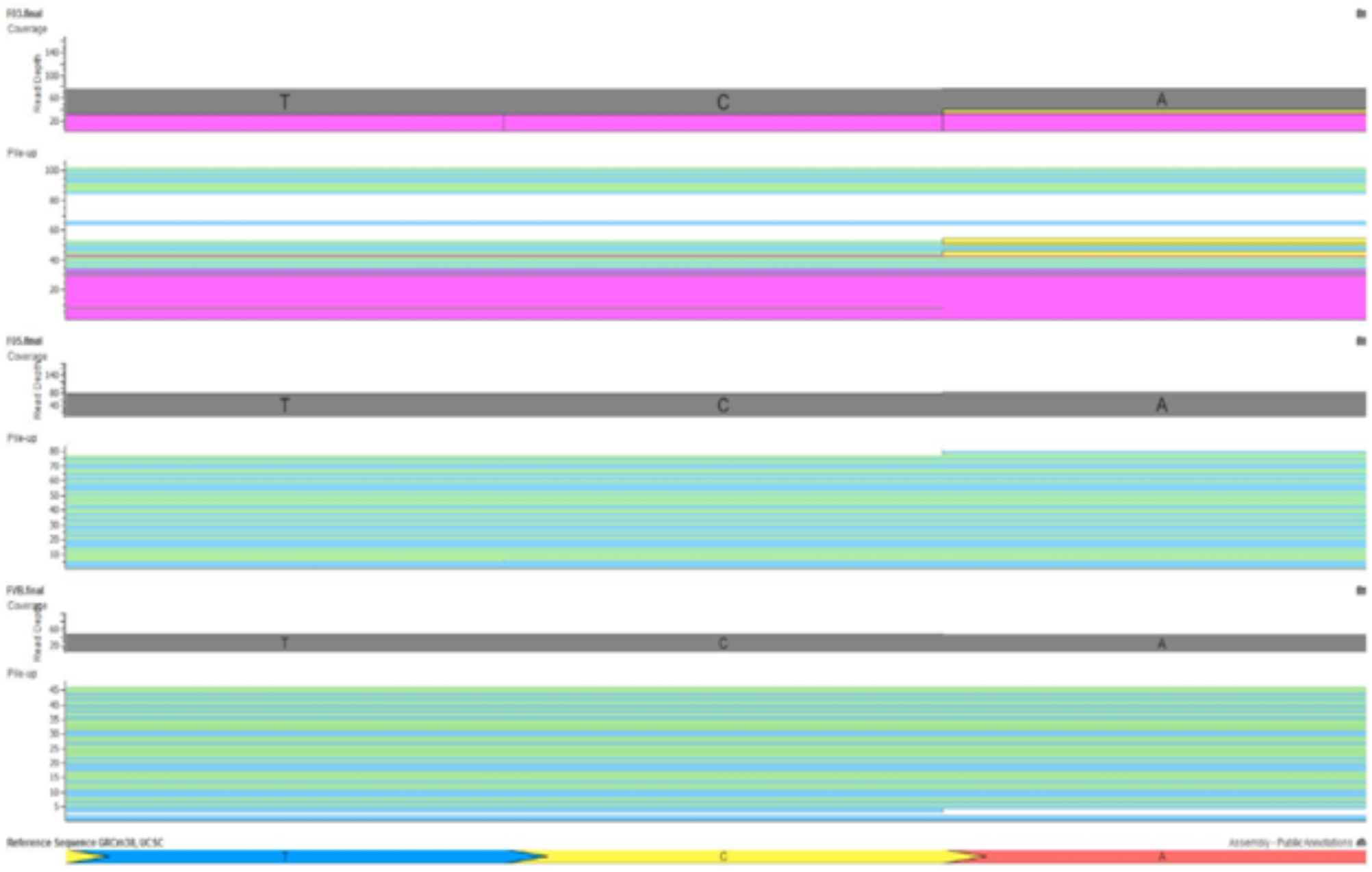
Whole genome sequence reveals more than 2 alleles at the chr1:86,357,000 – 86,357,003 deletion suggesting a CRISPR/Cas9-induced pattern of mutations. The F_0_3 mouse has numerous indel calls (pink) as well as an A>C SNV (yellow), while the F_0_5 and FVB control mice do not have multiple variants at this site (only wild-type). Images created using mouse BAM files and VarSeq (Golden Helix, Inc., Bozeman, Montana).

**Supplemental Figure 2.**
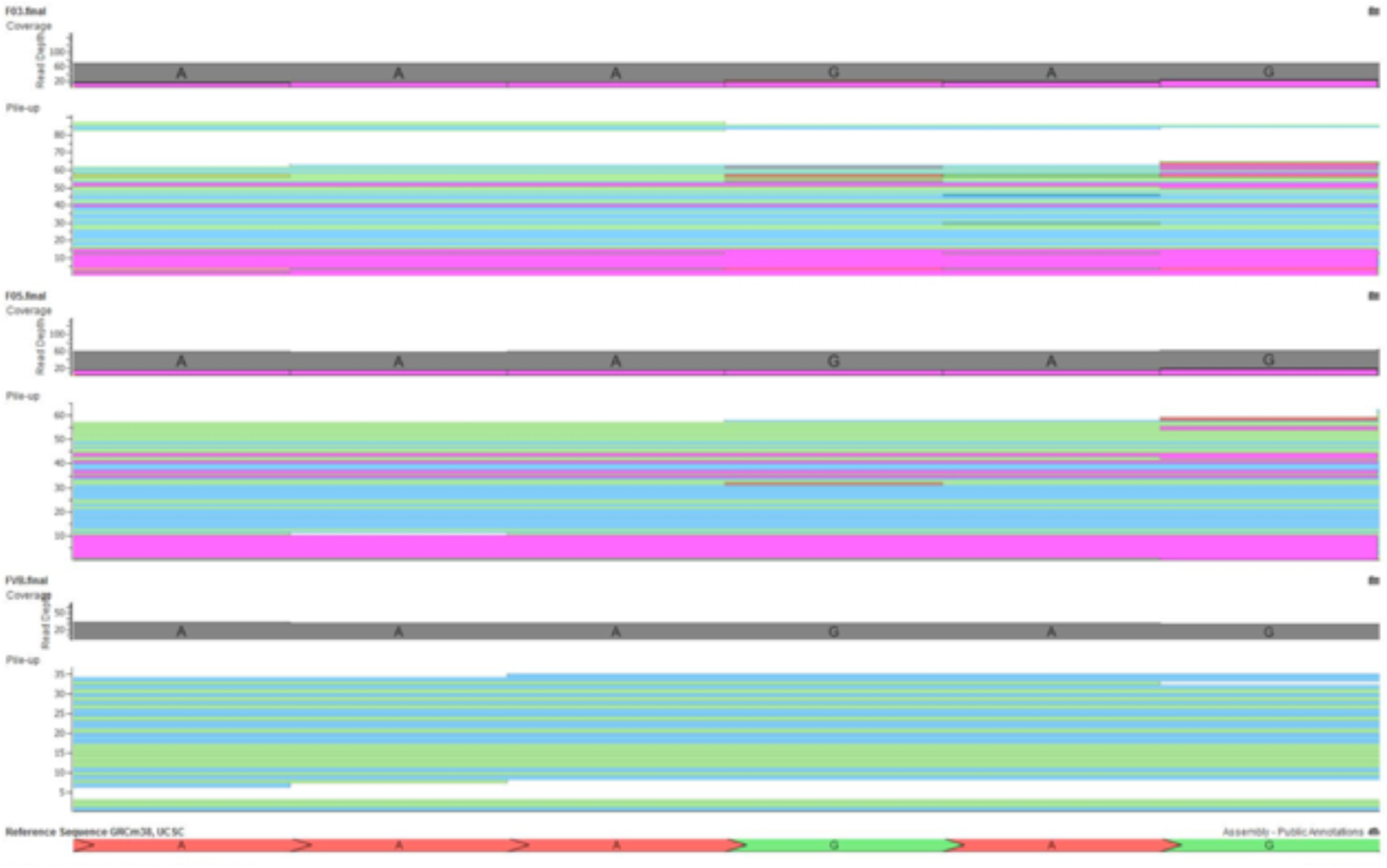
Whole genome sequencing reveals more than 2 alleles at the chr3:88,647,832 – 88,647,838 deletion suggesting a CRISPR/Cas9-induced pattern of mutations. F_0_3 and F_0_5 both have more than 2 alleles at the deletion site while FVB has only the wild-type allele at this site. Images created using mouse BAM files and VarSeq (Golden Helix, Inc., Bozeman, Montana).

**Supplemental Figure 3:**
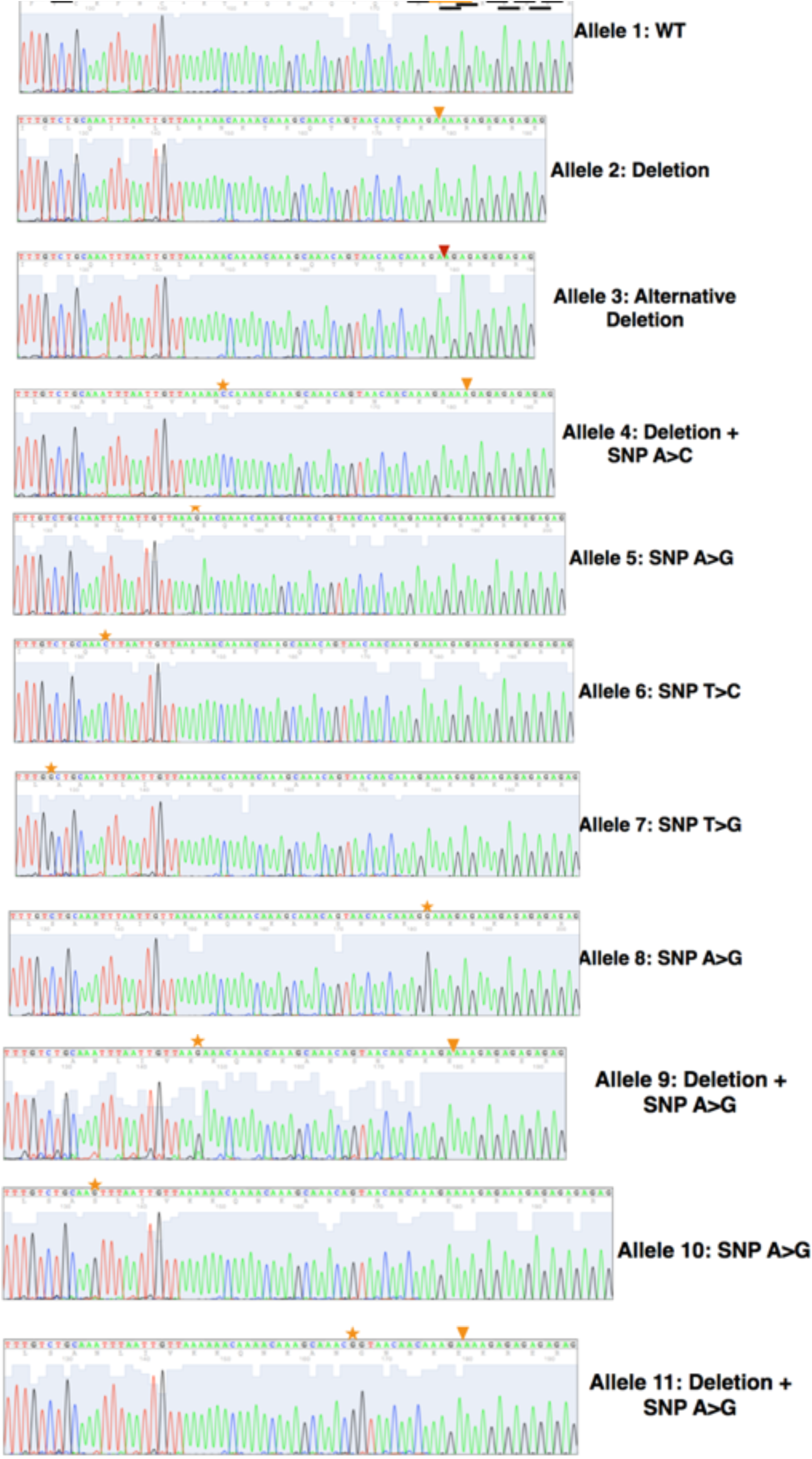
Sanger sequencing reveals more than 2 alleles at the chr3:88,647,832 deletion, suggesting a CRISPR/Cas9-induced pattern of mutations adjacent to NGG and NGA, which are underlined in black on the wild-type allele. Deletion site is underlined in orange on the wild-type allele and is noted by the orange arrow in alternative alleles. Alternative deletion is noted by the red arrow. Different SNVs are denoted by stars.

**Supplemental Figure 4.**
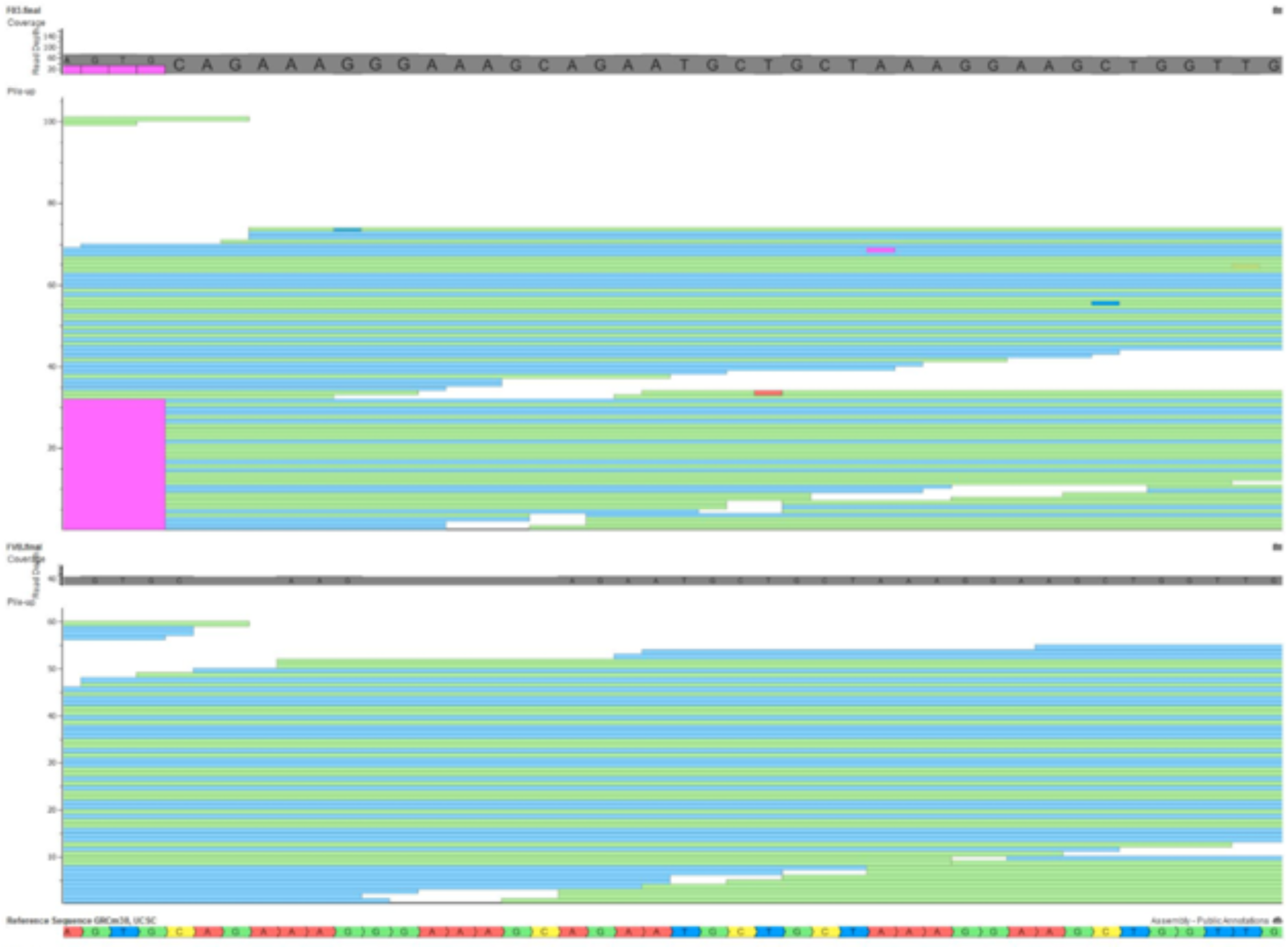
Whole genome sequencing reveals more than 2 alleles at the chr4:66,453,495 – 66,453,539 deletion suggesting a CRISPR/Cas9-induced pattern of mutations. F_0_3 has more than 2 alleles at the deletion site while FVB only has the wild-type allele at this site. Note that the right-most T>C SNV in F_0_3 was subsequently validated by Sanger sequencing (see **Figure 5**). Images were created using mouse BAM files and VarSeq (Golden Helix, Inc).

**Supplemental Figure 5.**
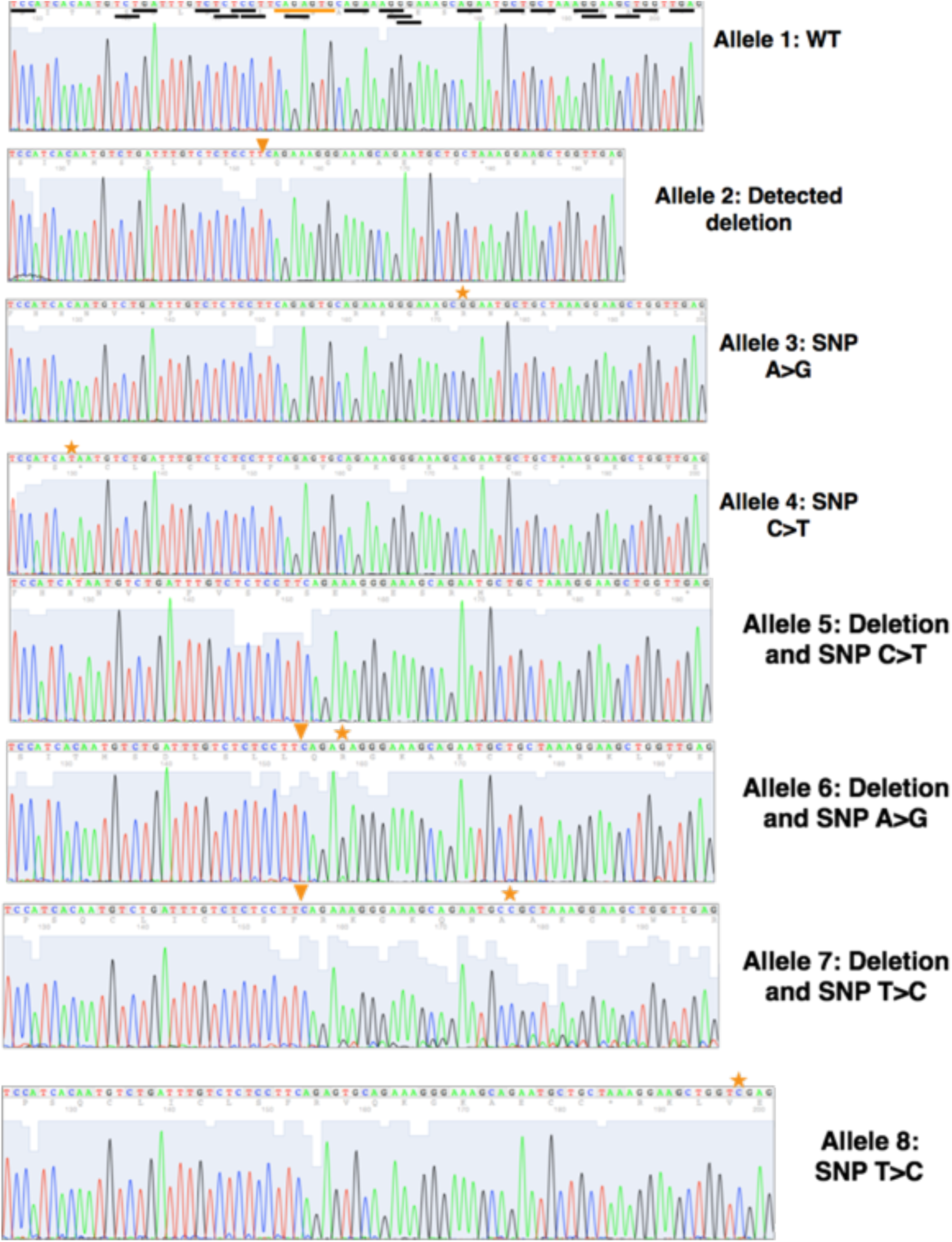
Sanger sequencing reveals more than 2 alleles at the chr4:66,453,495 deletion suggesting a CRISPR/Cas9-induced pattern of mutations adjacent to NGG and NGA which are underlined in black on the wild-type allele. Deletion site is underlined in orange on the wild-type allele and is noted by the orange arrow in alternative alleles. Different SNVs are denoted by stars. Note that the SNV in allele 8 was also present in the WGS data (see **Figure 4**).

**Supplemental Figure 6.**
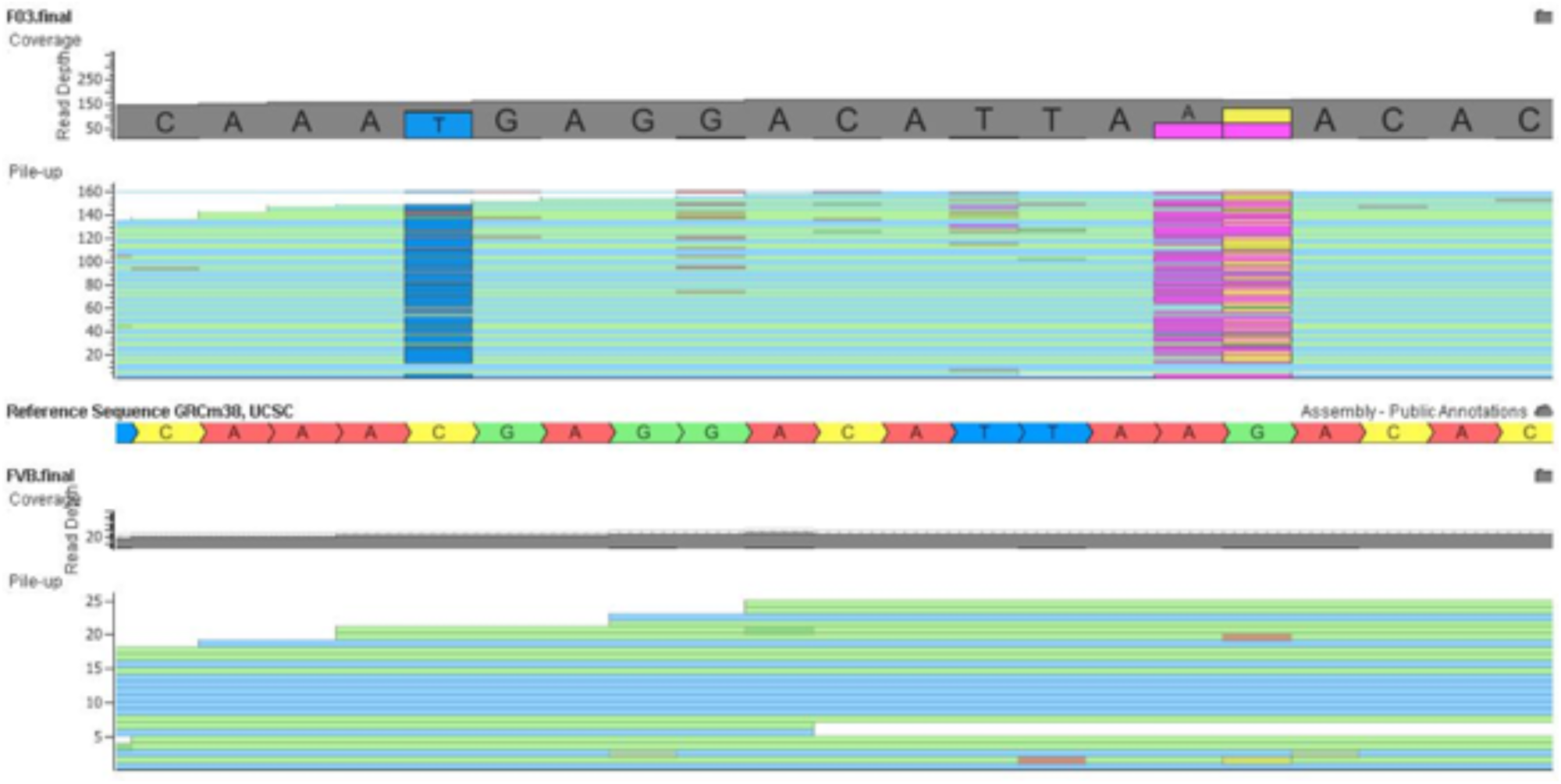
Whole genome sequencing reveals more than 2 alleles at the chrX:123,734,750-123,734,770 deletion, suggesting a CRISPR/Cas9-induced pattern of mutations. F_0_3 has more than 2 alleles at the deletion site while FVB only has the wild-type allele at this site. Images were created using mouse BAM files and VarSeq (Golden Helix, Inc). Note that one C>T and A>G SNV in F_0_3 was subsequently validated by Sanger sequencing (see **Figure 7**).

**Supplemental Figure 7.**
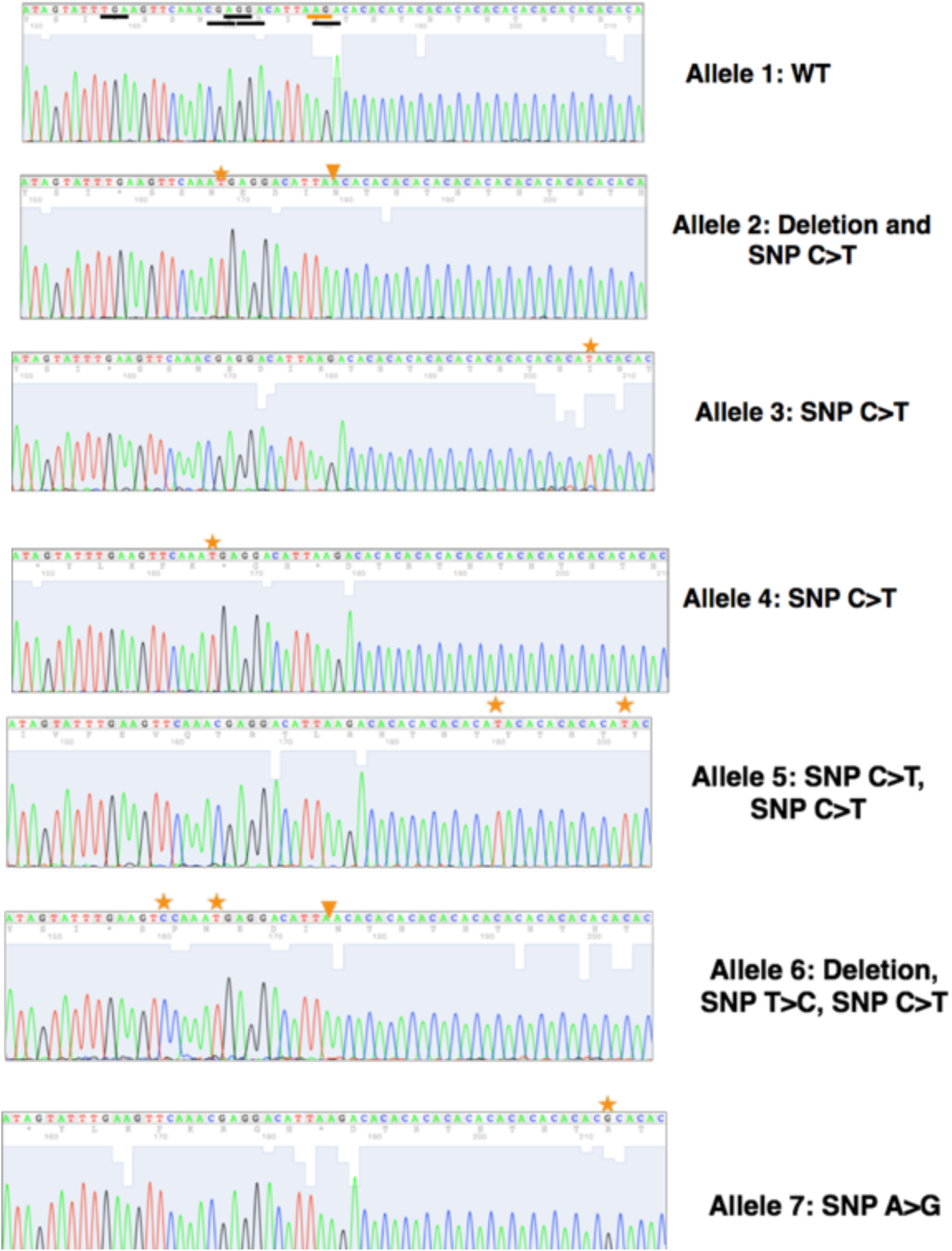
Sanger sequencing reveals more than 2 alleles at the chrX:123,734,765 deletion suggesting a CRISPR/Cas9-induced pattern of mutations adjacent to NGG and NGA which are underlined in black on the wild-type allele. Deletion site is underlined in orange on the wild-type allele and is noted by the orange arrow in alternative alleles. Different SNVs are denoted by stars. Note that the SNVs in allele 2 and 7 were also present in the WGS data (see **Figure 6**).

**Supplemental Figure 8:**
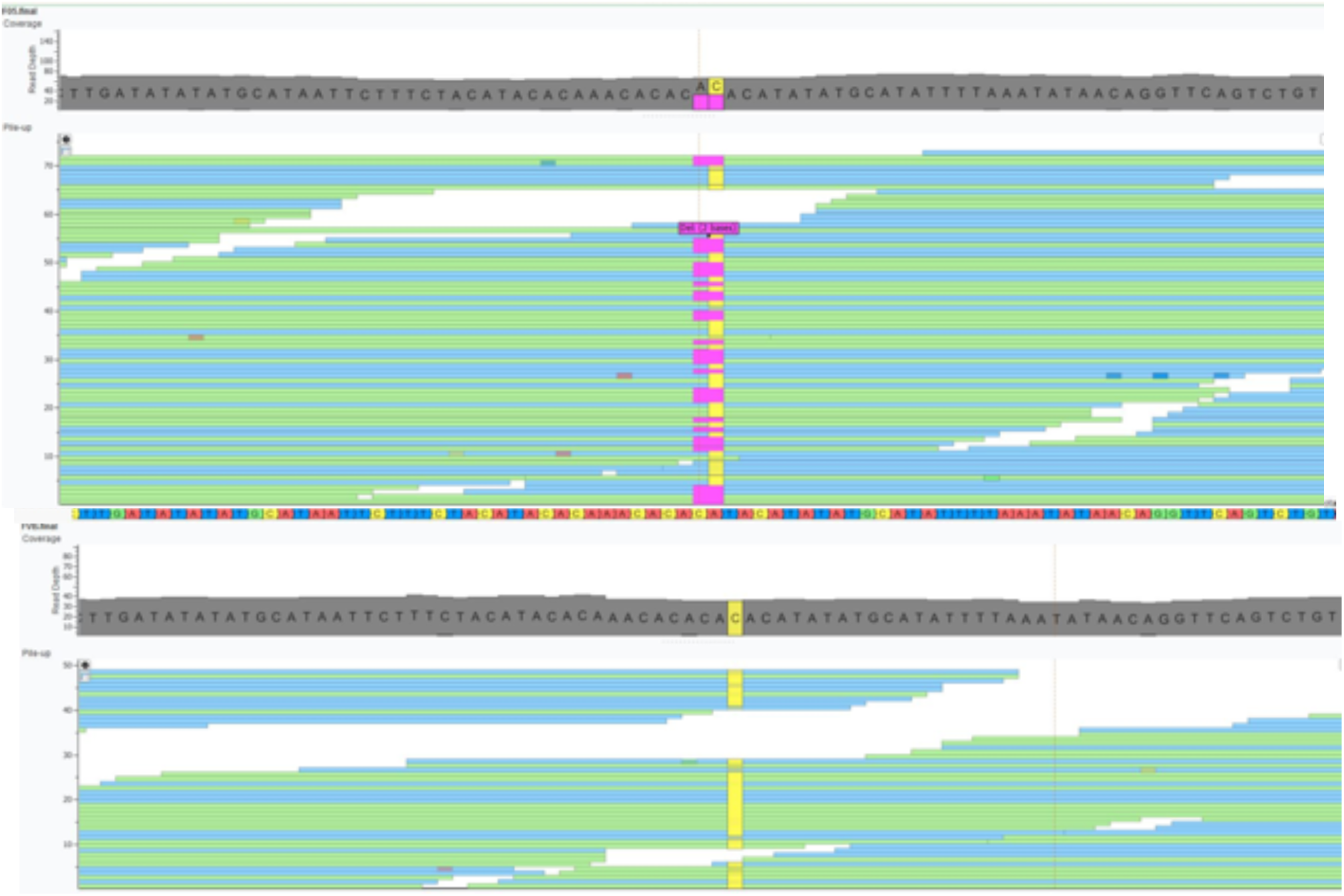
Whole genome sequencing reveals more than 2 alleles at the chr14:94,681,330-94,681,411 deletion suggesting a CRISPR/Cas9-induced pattern of mutations. F_0_5 has more than 2 alleles at the deletion site, while FVB only has the wild-type allele at this site. Images were created using mouse BAM files and VarSeq (Golden Helix, Inc). Note that a C>T SNV in F_0_5 was subsequently validated by Sanger sequencing (see **Figure 9**).

**Supplemental Figure 9.**
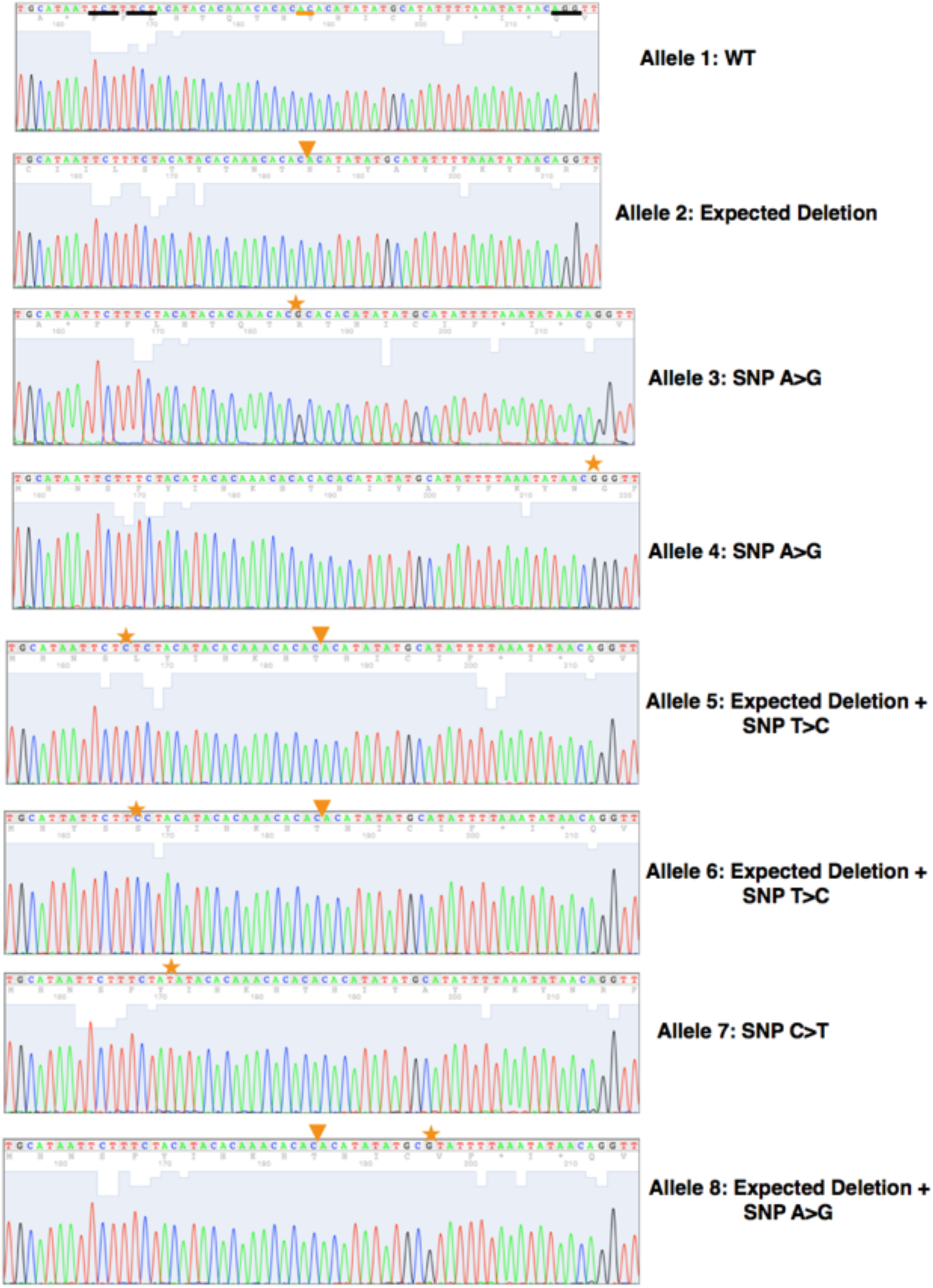
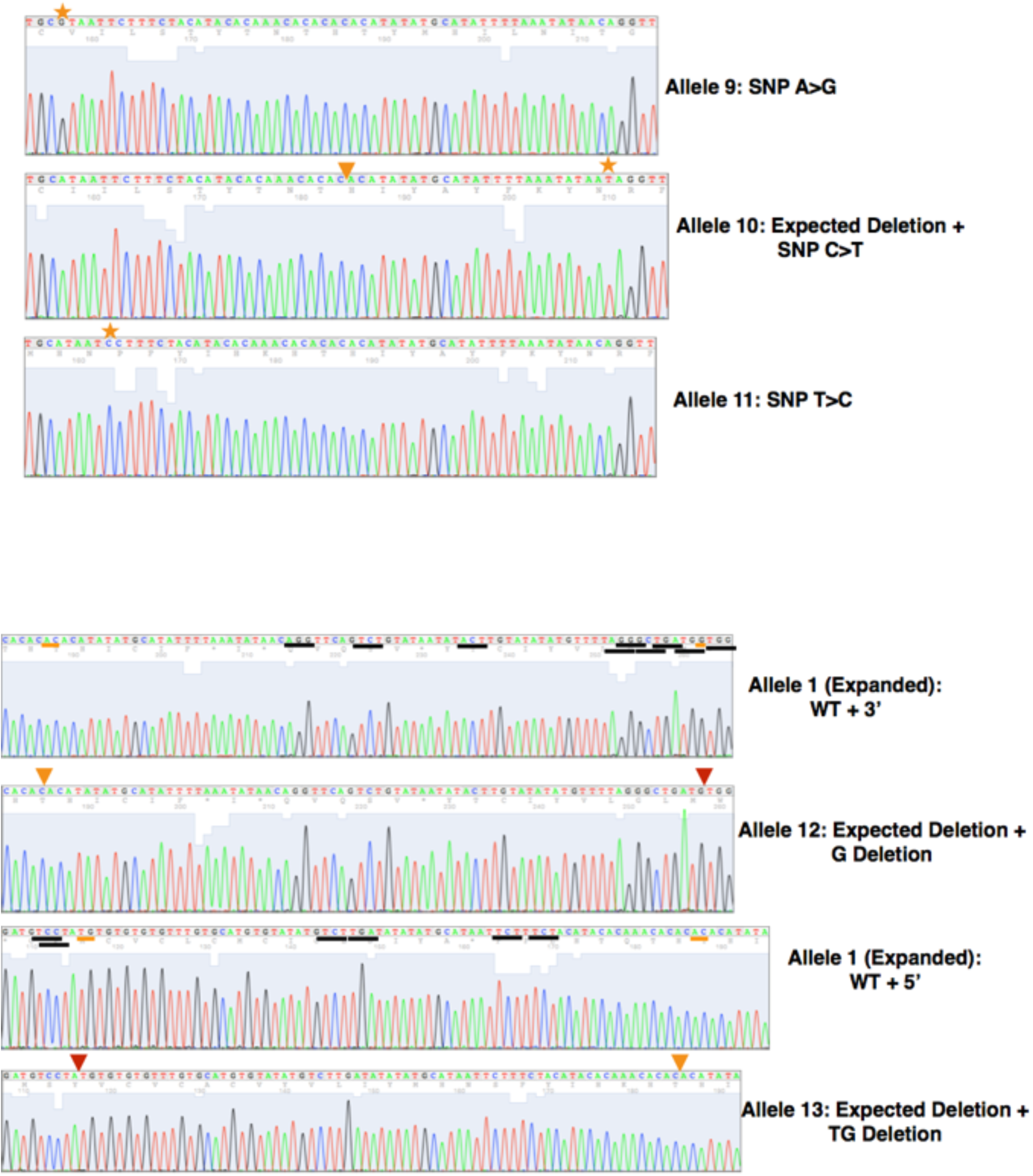
Sanger sequencing reveals more than 2 alleles, including SNVs and deletions, at the chr14:94681371 deletion, suggesting a CRISPR/Cas9-induced pattern of mutations adjacent to NGG and NGA which are underlined in black on the wild-type allele. Deletion site is underlined in orange on the wild-type allele and is noted by the orange arrow in alternative alleles. Additional deletions are noted by red arrows. Different SNVs are denoted by stars. Note that the C>T SNVs in allele 10 was also present in the WGS data (see **Figure 8**).

**Supplemental Figure 10.**
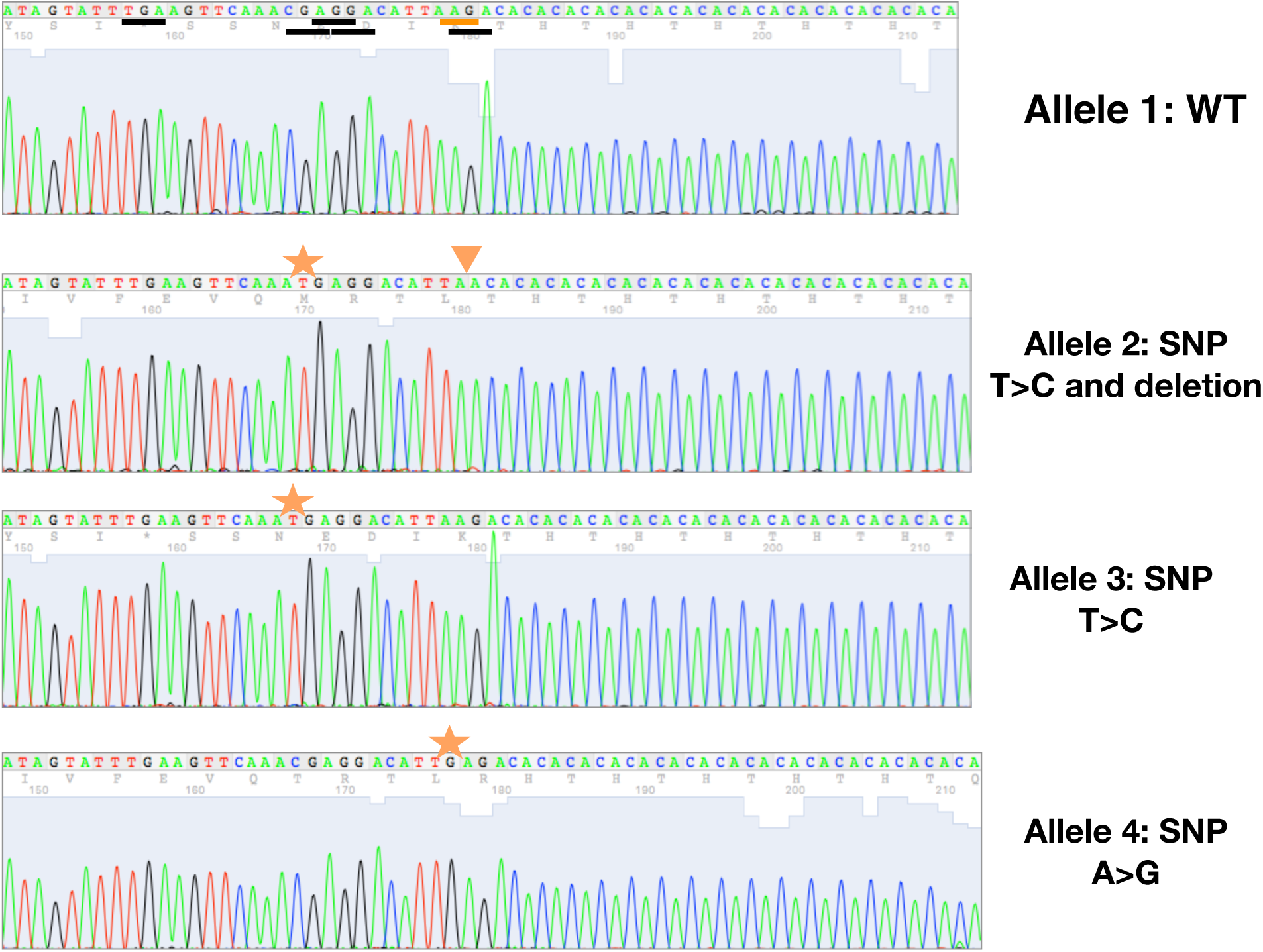
High-fidelity taq polymerase, topo cloning, and Sanger sequencing reveals more than 2 alleles at the chrX:123,734,765 deletion suggesting a CRISPR/Cas9-induced pattern of mutations adjacent to NGG and NGA, which are underlined in black on the wild-type allele. Deletion site is underlined in orange on the wild-type allele and is noted by the orange arrow in alternative alleles. Additional deletions are noted by red arrows. Different SNVs are denoted by stars. Note that allele 3 was also seen in WGS data (see **Figure 6**).

**Supplemental Figure 11.**
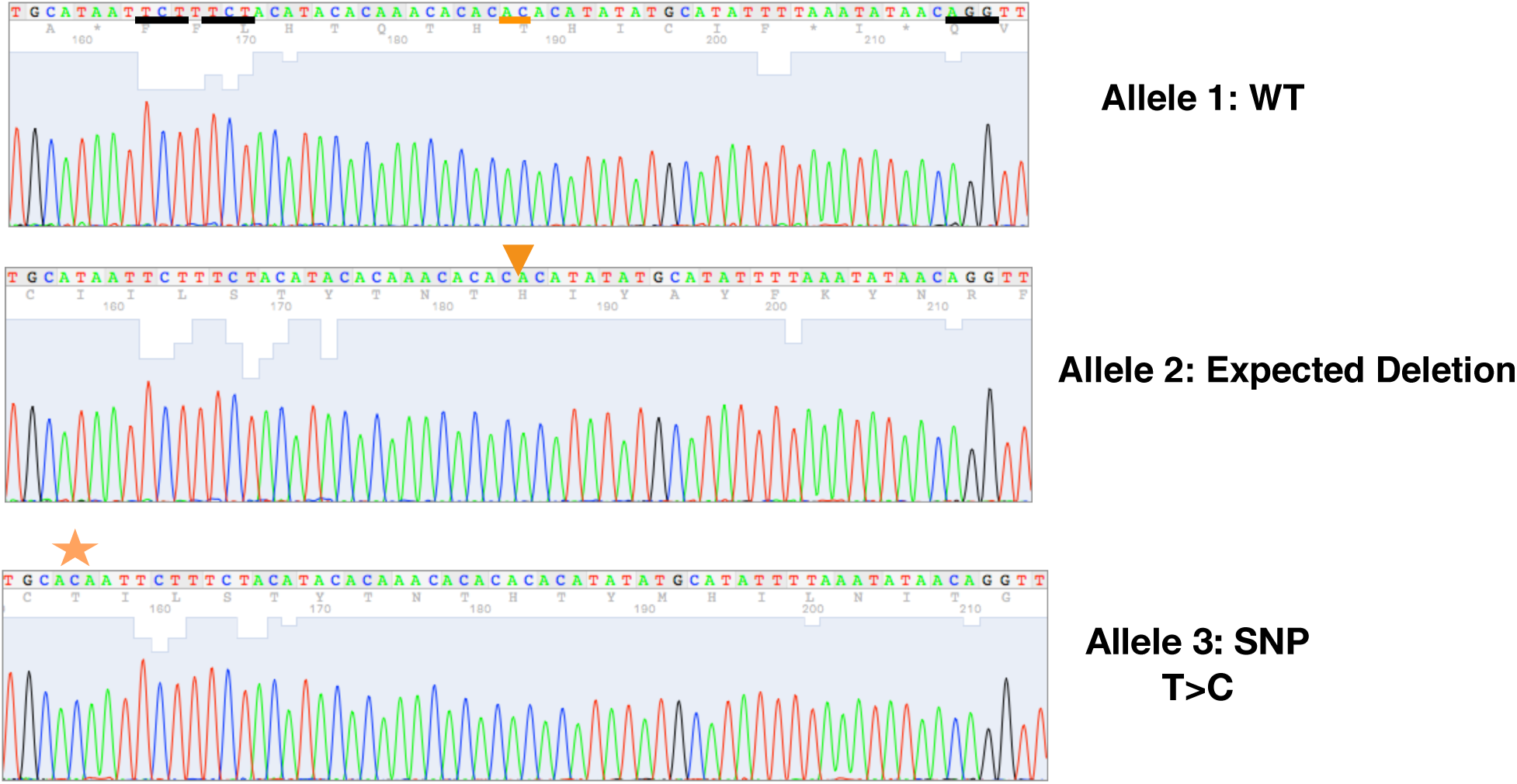
High fidelity taq polymerase topo cloning and sanger sequencing reveals more than 2 alleles at the chr14:94,681,317 deletion suggesting a CRISPR/Cas9-induced pattern of mutations adjacent to NGG and NGA which are underlined in black on the wild-type allele. Deletion site is underlined in orange on the wild-type allele and is noted by the orange arrow in alternative alleles. Additional deletions are noted by red arrows. Different SNVs are denoted by stars.

**Supplemental Figure 12.**
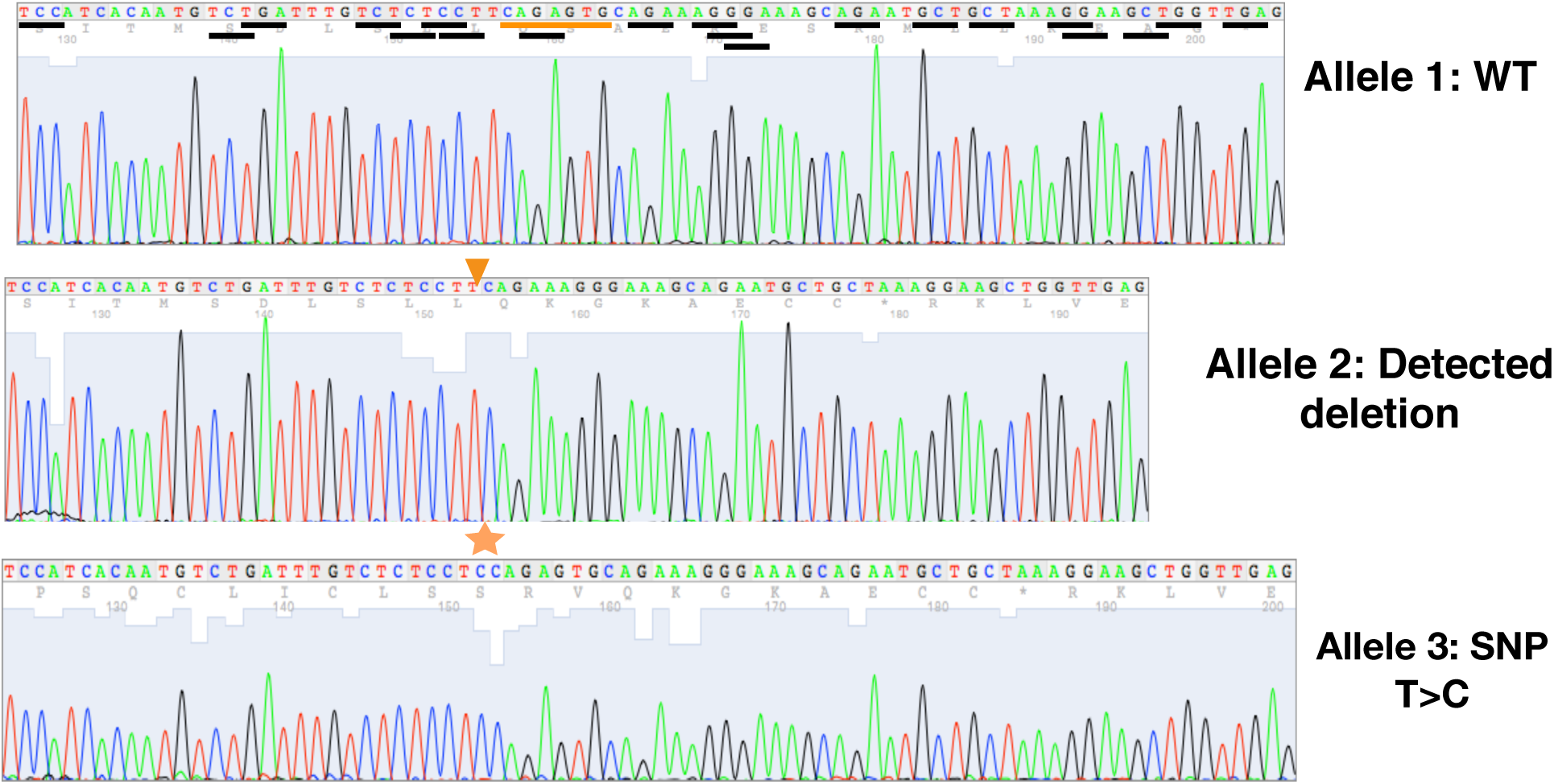
High fidelity taq polymerase topo cloning and Sanger sequencing reveals more than 2 alleles at the chr4:66,453,492 deletion, suggesting a CRISPR/Cas9-induced pattern of mutations adjacent to NGG and NGA which are underlined in black on the wild-type allele. Deletion site is underlined in orange on the wild-type allele and is noted by the orange arrow in alternative alleles. Additional deletions are noted by red arrows. Different SNVs are denoted by stars.

**Revised Supplemental Figure 3:**
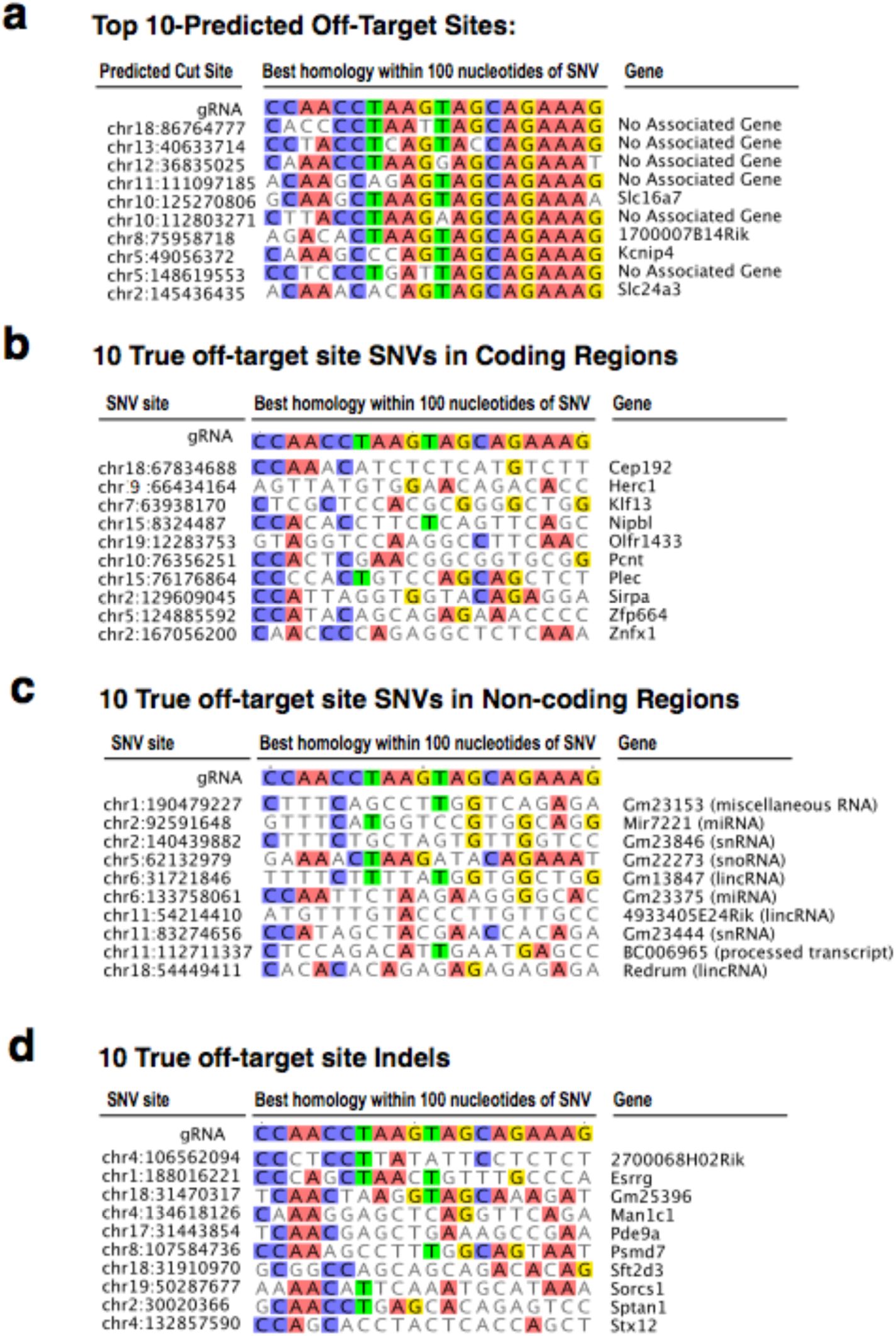
Sequence alignment of guide RNA to actual off-target regions does not show significant homology. **a**. The top-ten, off-target regions predicted *in silico*, by Benchling, to align to the gRNA. Sequences are 80 to 95% homologous to the gRNA. **b**. Regions surrounding ten, randomly selected, experimentally observed SNVs in coding regions aligned to the gRNA. Sequences are 15 to 45% homologous to the gRNA. **c**. Regions surrounding ten, randomly selected experimentally observed SNVs in non-coding region aligned to the gRNA. Sequences are 5 to 65% homologous to the gRNA. **d**. Regions surrounding ten randomly selected experimentally observed indels in both coding and non-coding region aligned to the gRNA. Sequences are 25 to 65% homologous to the gRNA.

